# The effect of the stress hormone cortisol on the metatranscriptome of the oral microbiome

**DOI:** 10.1101/339325

**Authors:** Ana E. Duran-Pinedo, Jose Solbiati, Jorge Frias-Lopez

## Abstract

Imbalances of the microbiome also referred to as microbial dysbiosis, could lead to a series of different diseases. One factor that has been shown to lead to dysbiosis of the microbiome is exposure to psychological stressors. Throughout evolution microorganisms of the human microbiome have developed systems for sensing host-associated signals such as hormones associated with those stressors, enabling them to recognize essential changes in its environment thus changing its expression gene profile to fit the needs of the new environment. The most widely accepted theory explaining the ability of hormones to affect the outcome of an infection involves the suppression of the immune system. Commensal microbiota is involved in stressor-induced immunomodulation, but other biological effects are not yet known. Here we present the impact that cortisol had on the community-wide transcriptome of the oral community. We used a metatranscriptomic approach to obtain first insights into the metabolic changes induced by this stress hormone as well as which members of the oral microbiome respond to the presence of cortisol in the environment. Our findings show that the stress hormone cortisol directly induces shifts in the gene expression profiles of the oral microbiome that reproduce results found in the profiles of expression of periodontal disease and its progression.

## INTRODUCTION

In recent years a considerable effort has been placed on characterizing the different microbial communities colonizing the human body ^1^. However, the nature of host-microbial interactions in the microbiome that allow for the maintenance of a stable microbiota is still poorly understood. Among the environmental factors that may alter the equilibrium in host-microbiome homeostasis, host-stress is a known risk factor for a variety of diseases. In case of acute stress, stress response may prepare the immune system for challenges such as infection, but when it becomes chronic, it may influence inflammatory processes leading to the development of systemic or local diseases such as rheumatoid arthritis ^2^, diabetes ^3^, or periodontitis ^4^. Furthermore, physiological stress can also alter the composition of the commensal microbiota in the human microbiome ^5^.

The most widely accepted theory to explain how hormones can influence microbial infections involves the of the immune system. According to this model, stress can activate the central nervous system and the hypothalamus releases corticotropin-releasing hormone and arginine vasopressin that stimulates the release of adrenocorticotropin from the pituitary, which in turn results in the production of cortisol by the adrenal cortex. However, almost immediately following its first use, cases of adrenaline-associated sepsis were reported ^6^. It was demonstrated that the dose of *Clostridium* needed to cause infection was significantly smaller when was injected in the presence of a therapeutic level of adrenaline ^7^. Since then, there have been reports associating the levels of neuroendocrine hormones, such as adrenaline, with infectious diseases, suggesting organisms themselves directly respond to the presence of stress hormones. The study of these interactions has been termed ‘microbial endocrinology’ ^8,9^. Microorganisms that have evolved systems for sensing host-associated signals such as hormones would have an evolutionary advantage over those that have not. Detecting such signals enables the microbiome to recognize essential changes in its environment thus changing its expression gene profile to fit the needs of the new environment. Moreover, there is also the possibility that the microbiome not only responds to human hormones but may be responsible for controlling their levels. Just recently it has been proposed that *Ruminococcus,* a member of the gut microbiome in piglets, controls levels of n-acetylaspartate, the second most abundant molecule in the brain, via levels of cortisol ^10^.

Although most investigations of stress hormones induction of growth and virulence have been carried out with gut-associated bacteria, a few studies have shown that stress hormones have a significant effect on the growth of periodontal pathogens ^11,12^. In the oral cavity glucocorticoids, including cortisol, depress immunity by inhibiting the production of secretory immunoglobulins, and neutrophil functions, all of which may impair defense against infection by periodontal microorganisms ^13^. Cortisol or hydrocortisone is the primary hormone responsible for the stress response, and its levels increase in saliva and serum with the severity of periodontal disease ^14,15^. We hypothesize that the oral microbiome is capable of sensing changes in the levels of stress hormones and its response could be associated with severity of periodontal disease. Here we present the results of an *in vitro* study to assess the effect that cortisol had on the community-wide transcriptome of the oral community. We used a metatranscriptomic approach to obtain first insights into the metabolic changes induced by this stress hormone as well as which members of the oral microbiome respond to the presence of cortisol in the environment.

## RESULTS

Our first experiments consisted of treating samples of dental plaque with cortisol at a concentration found in the saliva of patients with periodontitis ^14,16^. As a control for these experiments we used dental plaque that was incubated under the same conditions as the treatment samples but without cortisol added to the saliva used as the medium. People with periodontal disease present higher levels of cortisol in the gingival crevicular fluid ^14^, a serum exudate in direct contact with the oral microbiome. After only 2 hours of incubation in the presence of cortisol, we proceed to perform the analysis, avoiding possible changes related to the growth of individual members of the microbiome and not to the presence of the hormone itself (see Methods section in the Supplementary Information accompanying this manuscript). We assigned the phylogenetic origin of those sequences using Kraken^17^, and phylogenetic profiles were used to identify significant differences between active communities under the different conditions studied by performing linear discriminant analysis (LDA) effect size (LEfSe). In our study, LEfSe determines the active members of the oral microbiome, based on changes in the number of transcripts, that most likely explain differences between the two biological conditions tested (presence and absence of cortisol). LEfSe robustly identifies features that are statistically different among biological classes using the non-parametric factorial Kruskal-Wallis (KW) sum-rank test and subsequently uses a set of pairwise tests among subclasses using the (unpaired) Wilcoxon rank-sum test for biological consistency. Finally, LDA estimates the effect size of each differentially abundant taxonomic group ^18^. Among all the organisms in the oral community, members of the phylum Fusobacteria (class Fusobacteriia and order Fusobacteriales) were significantly more active (increased the number of transcripts significantly) after the addition of cortisol (Fig. 1). Among them, one species, *Leptotrichia goodfellowii,* was substantially more active (Fig. 1b). Species belonging to the *Fusobacteriales*, such as *Fusobacterium nucleatum* have been associated with a wide variety of human diseases, other than periodontitis, including adverse pregnancy outcome, GI disorders (e.g., colorectal cancer, inflammatory bowel disease), cardiovascular disease, rheumatoid arthritis, and respiratory tract infections ^19^. *Leptotrichia* species are typically part of the commensal flora in the oral cavity and genitourinary tract and are seldom found in clinically significant specimens. However, *Leptotrichia* has been found to be in higher proportion in gingivitis ^20,21^.

**Figure 1.**
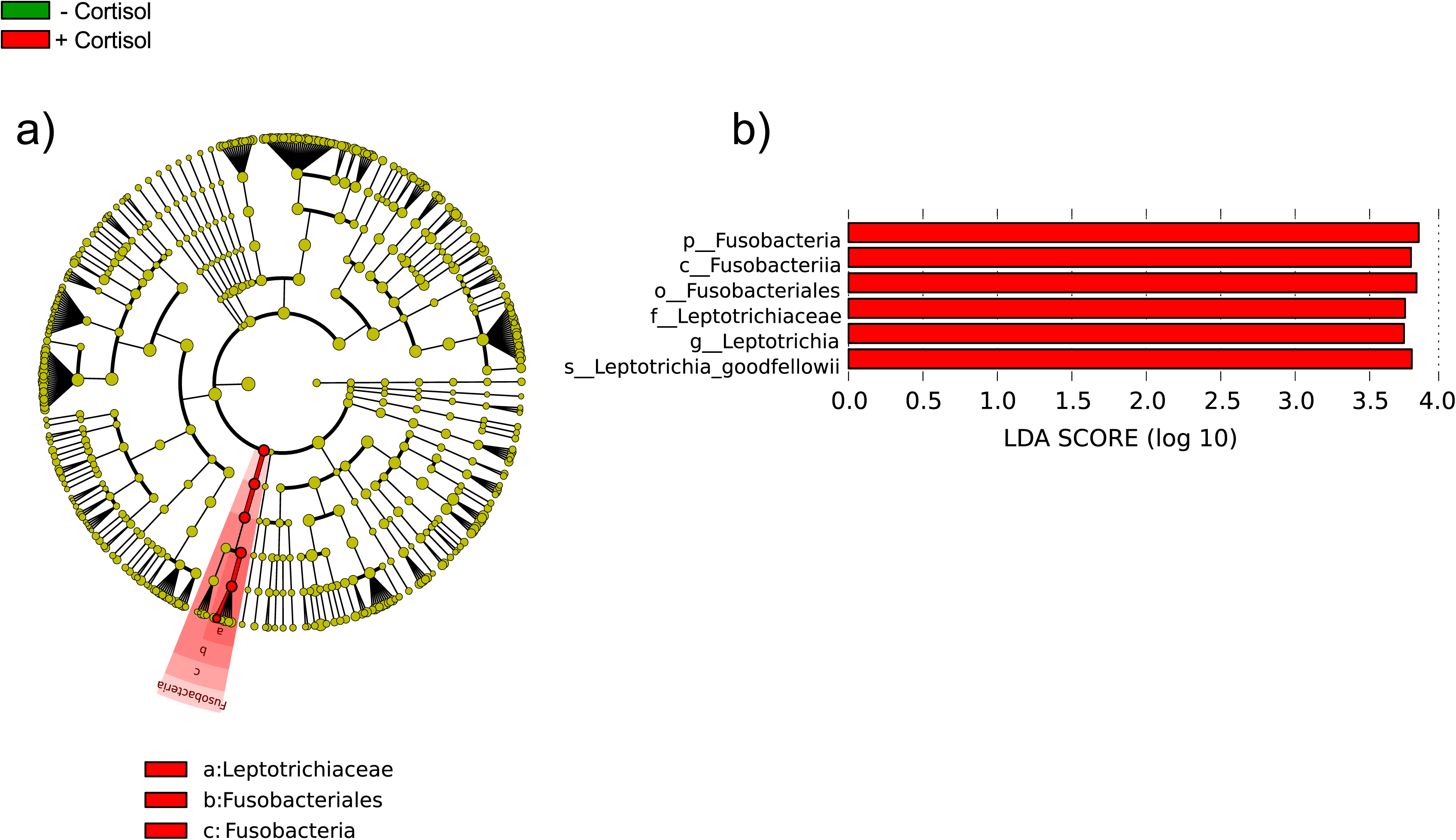
Distinct active taxa identified using LEfSe analysis. Metatranscriptome hit counts were obtained using Kraken against an oral microbiome database. Counts were then analyzed using LEfSe to identify significant differences at the species level between the microbial communities compared. a) Cladogram constructed using the LEfSe method to indicate the phylogenetic distribution of active bacteria that were remarkably enriched. Red represented the enriched taxa the untreated microbial community and green represented the enriched taxa in the microbial community treated with cortisol. b) LDA (Linear Discriminant Analysis) scores showed significant bacterial differences within groups at different taxonomic levels. Red represented the enriched taxa in the untreated microbial community and green represented the enriched taxa in the microbial community treated with cortisol.

Next, we looked at how cortisol influenced the profiles of expression of the oral microbiome. To this end, we performed enrichment of Gene Ontology (GO) terms analysis, and we observed that, after only two hours of exposure to the hormone, the profiles of activities of the whole community had already changed and they were similar to the profiles of expression we previously found in periodontitis progression ^22,23^. Among them GO terms associated with proteolysis, oligopeptide transport, iron metabolism, and flagellum assembly were over-represented in the presence of cortisol (Fig. 2a). What is more important these activities have been associated with an increase in pocket depth, and essential clinical parameter in periodontitis, during the progression of the disease ^23^.

**Figure 2.**
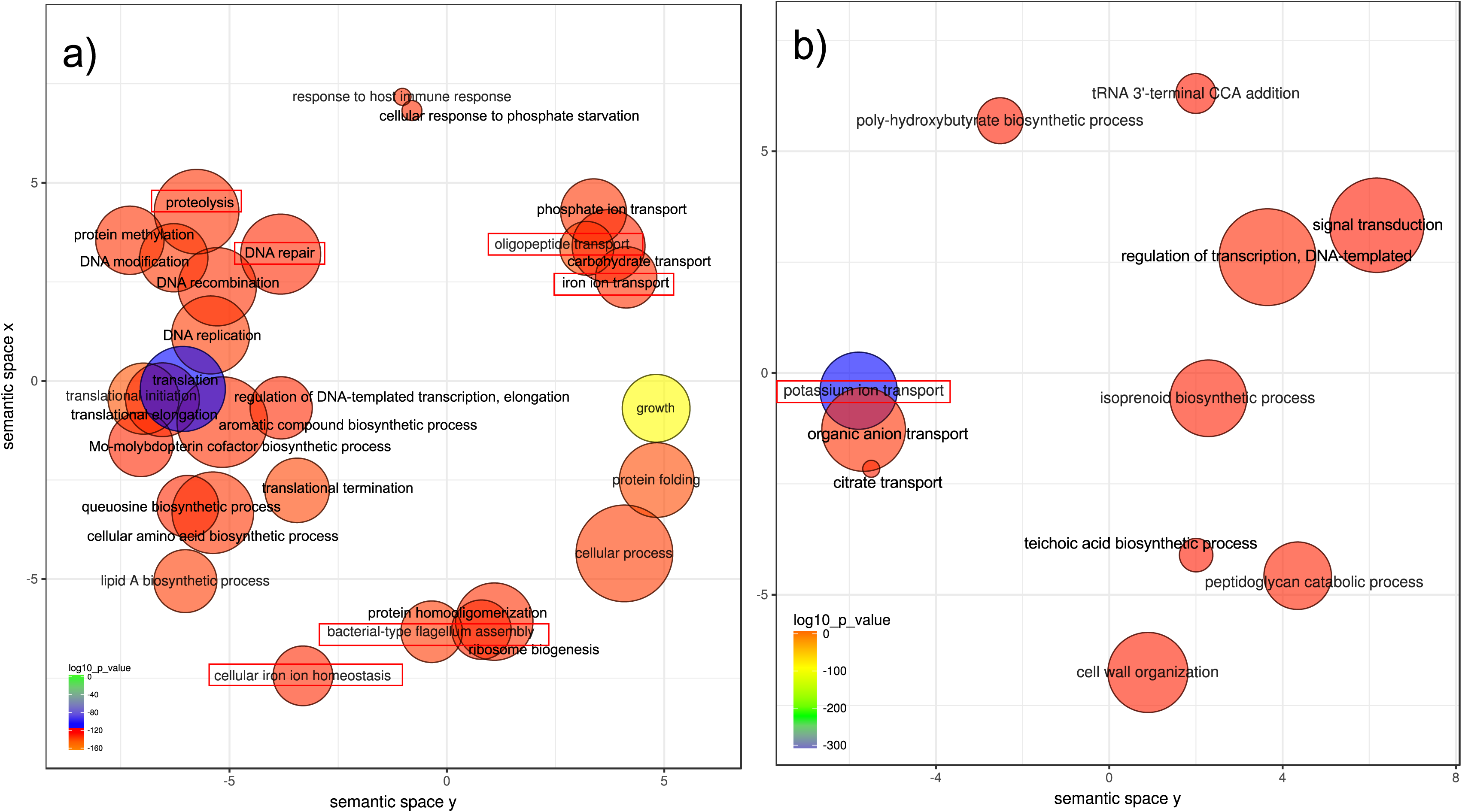
Changes in functional profiles in the oral microbiome treated with cortisol. We used Gene Ontology (GO) enrichment analysis to assess the oral microbiome functional response to the presence of added cortisol to the medium. Enriched terms obtained using GOseq were summarized and visualized as a scatter plot using REVIGO. Only GO terms with FDR adjusted p-value < 0.05 in the ‘GOseq’ analysis were used. a) Over-represented functional activities in the presence of cortisol summarized as GO terms related to biological processes. b) Over-represented functional activities in the absence of cortisol summarized as GO terms related to biological processes. Circle size is proportional to the frequency of the GO terms; color indicates the log10 p-value (red higher, blue lower). The distance between circles represents GO terms’ semantic similarities. Each of the circles represents a GO term, which depending on the similarity in the terms included in them they will be closer or more distant in the graph. In red are activities we have previously seen associated with periodontitis and its progression ^22,23^.

Interestingly, GO terms linked to response to host immune response were also over-represented when cortisol was present, even though the microbiome in our samples was not in contact with host cells (Fig. 2a). As previously indicated potassium ion transport was significantly under-represented when cortisol was added (Fig. 2b), mimicking our *in vivo* observations of periodontitis progression ^23^. In a follow-up manuscript, we demonstrated that ion potassium is a signal in functional dysbiosis of periodontal disease ^24^.

Interestingly, when we look at the activities associated explicitly with putative virulence factors, we found a similar pattern of over-represented GO terms related to the addition of cortisol (Sup. Fig 1). Although phylum Fusobacteria was the phylogenetic group that increased its transcriptional activity more significantly, the rest of the microbiome showed a shift in its profile of expression. Thus, the more significant fraction of up-regulated putative virulence factors seems to be synthesized by members of the genus *Streptococcus* (Sup. Fig 2). We observed similar results in our two previous studies on periodontitis progression ^23^ and the effect of ion potassium in functional dysbiosis of the oral microbiome ^24^. As a whole, these results seem to indicate that the presence of cortisol leads to a community-wide response very similar to the one observed *in vivo* during periodontitis.

We then used a simpler system based on our previous results on the community-wide transcriptome results. To this end, we performed transcriptome analysis of the effect of cortisol on pure culture of organisms that we identified as more active in the presence of cortisol (Fig. 1): *Leptotrichia goodfellowii* and *Fusobacterium nucleatum*, which is a representative of the order Fusobacteriales and an essential member of the oral microbiome. In just 2 hours of exposure, there was a shift in their transcriptome profiles. *F. nucleatum* showed an increase in biological processes GO terms associated with proteolysis, cobalamin biosynthesis and iron transport (Sup. Fig 3a), which we have previously found associated with the progression of periodontal disease ^22,23^. In the case of *L. goodfellowii,* we also found activities related to iron ion transport (Sup. Fig 3b). The intersection of metabolic events that were commonly altered in *F. nucleatum* and *L. goodfellowii* was associated with the growth of the organisms such as lipid A biosynthesis, DNA replication or translation (Sup. Fig 3), which indicates activation of their metabolism.

Moreover, iron transport was also altered in both organisms in the presence of cortisol (Sup. Fig 3). These results are in agreement with the effects observed for the whole oral microbiome where members of the Fusobacteriales order are more active when cortisol was added to the medium, (Fig. 1). Likewise, the intersection of common molecular functions enriched with the addition of cortisol in *F. nucleatum* and *L. goodfellowii* include iron acquisition (iron-ion binding and iron-ion transmembrane transporter activity) and peptidase activities (serine-type endopeptidase activity) (Sup. Fig 4).

Altogether, these results show that the only exposure to cortisol in the oral microbiome is enough to cause a significant shift in the gene expression profile of the community mimicking community-wide expression profiles observed in periodontitis and its progression *in vivo* ^22,23^. Our results also highlight the importance of *Fusobacteria* and *Leptotrichia* as the members of the community that more rapidly increase their metabolism in response to an increase of cortisol in the environment.

Though it has been known for a long time that hormones had some effect on the metabolism of bacteria ^7,25–27^, the mechanisms by which this cross-talk happens to remain mostly unknown. As we deepen our understanding of the precise roles of the microbiome in health and disease, we expect that new mechanisms will be shown to involve host hormones, including novel interactions. The present manuscript builds on the idea that human hormones can be used by the microbiome as signals to sense changes in their environment thus modifying its expression profile to fit the new conditions better. Nonetheless, we should recognize that given the limited number of samples in this pilot study, further work with a larger sample size should be performed to confirm the results presented in this manuscript.

Using a metatranscriptomic analysis would allow us to assess the direct effect that hormones have on the metabolism of the microbial community. Approaching the host-microbiome interactions from a microbial endocrinology-based point may provide an understanding of the specific pathways by which microorganisms may influence the outcome of certain infections and chronic diseases.

## Data Availability

The sequence datasets used in these analyses were deposited at the Human Oral Microbiome Database (HOMD) under the submission number 20180522 (ftp://homd.org/publication_data/20180522/).

## ACKNOWLEDGMENTS

This research reported was supported by the National Institute of Dental and Craniofacial Research of the National Institutes of Health (NIDCR/NIH) under award number DE021553 and DE021553-05A1.

## AUTHOR CONTRIBUTIONS

JFL and ADP designed the experiments. ADP and JS performed the experiments. JFL, ADP, and JS wrote the manuscript.

## Supporting Information Legends

**Supplementary Figure 1. Community-wide GO enrichment analysis of differentially expressed virulence factors in response to the presence and absence of added cortisol to the medium.** Putative virulence factors were identified by alignment of the protein sequences from the different genomes against the Virulence Factors Database (VFDB) as described in the methods section. Enriched terms obtained using GOseq were summarized and visualized as a scatter plot using REVIGO. Only GO terms with FDR adjusted p-value < 0.05 in the ‘GOseq’ analysis were used. A) Summarized GO terms related to biological processes after addition of cortisol. B) Summarized GO terms related to biological processes with no cortisol added. Circle size is proportional to the frequency of the GO terms; color indicates the log10 p-value (red higher, blue lower). The distance between circles represents GO terms’ semantic similarities. Each of the circles represents a GO term, which depending on the similarity in the terms included in them they will be closer or more distant in the graph. In red are activities we have previously seen associated with periodontitis and its progression ^22,23^.

**Supplementary Figure 2. Species up-regulating putative virulence factors in the presence of cortisol.** We ranked the species based on the number of up-regulated putative virulence factors observed in the metatranscriptome of the oral microbiome. Putative virulence factors were identified by alignment of the protein sequences from the different genomes against the Virulence Factors Database (VFDB) as described in the methods section. Numbers in the graph refer to the percentage of hits of the different species for the putative virulence factors identified. We selected only species whose percentage of putative virulence factors from the total of the community was higher than 1%. We included also results from 2 previous studies one on periodontal disease progression ^23^ and another where we showed that potassium was a crucial signal in dysbiosis ^24^. In red, species that were ranked at the top of putative virulence factors up-regulation in all three studies.

**Supplementary Figure 3. Changes in functional profiles in pure cultures of *Fusobacterium nucleatum* and *Leptotrichia goodfellowii*.** We used Gene Ontology (GO) enrichment analysis to assess the functional response of *Fusobacterium nucleatum* and *Leptotrichia goodfellowii* to the presence of added cortisol to the medium. Biological processes GO terms associated with changes in gene expression profiles in *Fusobacterium nucleatum* and *Leptotrichia goodfellowii*. GO terms were assigned to differentially expressed genes due to the addition of cortisol and summarized using REVIGO. a) GO terms associated with up-regulated genes in *F. nucleatum* b) GO terms associated with down-regulated genes *L. goodfellowii*. In green are metabolic activities that were associated with up-regulated genes in both *F. nucleatum* and *L. goodfellowii*. In red are activities we have previously seen associated with periodontitis ^22,23^.

**Supplementary Figure 4. Common molecular function GO terms associated with changes in gene expression profiles in *Fusobacterium nucleatum* and *Leptotrichia goodfellowii*.** Gene Ontology (GO) terms were assigned to differentially expressed genes due to the addition of cortisol and summarized using REVIGO. Networks of over-represented molecular functions from *F. nucleatum* and *L. goodfellowii* were then uploaded to Cytoscape, and the intersection of the two networks was extracted and plotted as shown in the figure. In red are activities we have previously seen associated with periodontitis ^22,23^.

## Supplementary methods

### Challenging the oral microbiome with cortisol

To assess the effect that cortisol has on the oral microbiome we performed metatranscriptome analysis of a biological triplicate of subgingival dental plaque from a periodontally healthy subject. The periodontal status of the donor was evaluated at the Forsyth Institute Clinic according to criteria described by the American Academy of Periodontology^1^ (presented ≤ 10% of sites with BOP, no PD or CAL > 3 mm). Samples were taken separately from six individual molar sites with no BOP and PD or CAL < 3 mm using sterile Gracey curettes. Plaque samples and saliva came from the same subject pooled and were resuspended in 6 ml of saliva. The saliva+plaque suspension was mixed by gentle vortexing and aliquoted in 1ml volume per well in a 24-well Corning™ Costar™ Flat Bottom Cell Culture Plate (Thermo Fisher).

To three of those wells, we added hydrocortisone-water soluble (Sigma-Aldrich) to a final cortisol concentration of 3.5μg/ml (10μl per well of a dilution 1/100 from a stock solution of 35mg/ml), while we did not add anything to the other three wells that were used as controls. The plate was incubated at 37° C for 2 hours under anaerobic conditions. Cells were collected by centrifugation at 10,000 x g for 5 minutes, and RNA was extracted immediately for further analysis as described below.

### Effect of cortisol on *Leptotrichia goodfellowii* and *Fusobacterium nucleatum*

We also challenged *L. goodfellowii* and *F. nucleatum* with cortisol to assess the effect that this hormone has on their expression profiles. Both organisms were grown on 10mL of BD BBL Schaedler broth (Thermo Fisher) at 37°C under anaerobic conditions until reaching an OD_600_ = 0.5. Cells were collected by centrifugation at 10,000 x g for 5 minutes, washed and resuspended in 10 ml of modified saliva medium (MSM) as described by Pratten et al. ^2^. We used artificial saliva medium to avoid variability among actual saliva samples from volunteers that could influence the results. MSM composition was: yeast extract 2 g/l (BD Biosciences), proteose peptone 5 g/l (Sigma-Aldrich), porcine gastric mucin 2.5 g/l (Sigma-Aldrich), sodium chloride 0.2 g/l, potassium chloride 0.2 g/l (Sigma-Aldrich), calcium chloride 0.3 g/l; 1.25 ml/l of a 0.2 μm filter-sterilized solution of 40% urea was added after autoclaving. MSM medium was allowed to stay at least 60 h under anaerobic conditions before inoculation.

1ml of the suspension per well was added to a 24-well Corning™ Costar™ Flat Bottom Cell Culture Plate (Thermo Fisher). To three of those wells, we added hydrocortisone/cortisol (Sigma-Aldrich) to a final cortisol concentration of 3.5μg/ml while three wells with the suspension of the microorganisms but without cortisol were used as controls. The plate was incubated at 37°C for 2 hours under anaerobic conditions. Cells were collected by centrifugation at 10,000 x g for 5 minutes, and RNA was extracted immediately for further analysis as described below. In the case of these pure cultures, RNA was not amplified for analysis. The rest of the procedure is identical to the one used for metatranscriptomic analysis of the whole community.

### Metatranscriptomic analysis

Detailed protocols for community RNA extraction, RNA amplification, and Illumina Sequencing are described in Yost et al. ^3^. Briefly, 600μL of mirVana kit lysis/binding buffer and 300 μl of 0.1-mm zirconia-silica beads (BioSpec Products) were added to the samples. Samples were bead beaten for 1 min at maximum speed. RNA was extracted following the protocol of *the mir*Vana™ Isolation kit for RNA. MICROB*Express* (Life Technologies) was used to remove prokaryotic rRNA. All kits were used following the manufacturer’s instructions. RNA amplification was performed on total bacterial RNA using MessageAmp ™ II-Bacteria RNA amplification kit (Life Technologies) following the manufacturer’s instructions. Sequencing was performed at the Forsyth Institute.

For the bioinformatic analysis, we first calculated the biological variation (BCV) after estimating the common dispersion using the R package edgeR. These values were used as a cv (coefficient of variation) cutoff in NOISeq. For the whole community, analysis cv was 2 (200 cutoff NOISeq), for the *Leptotrichia goodfellowii* libraries cv was 1.2 (120 cutoff NOISeq) and for the *Fusobacterium nucleatum* libraries cv was 0.28 (28 cutoff NOISeq).

Low-quality sequences were removed from the query files. Fast clipper and fastq quality filter from the Fastx-toolkit (http://hannonlab.cshl.edu/fastxtoolkit/) were used to remove short sequences with a quality score >20 in >80% of the sequence. Cleaned files were then aligned against the bacterial/archaeal database using bowtie2. We generated a .gff file to map hits to different regions in the genomes of our database. Read counts from the SAM files were obtained using bedtools multicov from bedtools ^4^.

To identify deferentially expressed (DE) genes from the RNA libraries, we applied non-parametric tests to the normalized counts using the NOISeqBio function of the R package ‘NOISeq’ with ‘tmm’ normalization, with batch and length correction and removing genes whose sum of hits across samples was lower than 10. We used a significance threshold value of q=0.95, which is equivalent to an FDR adjusted p-value of 0.05^5^.

To evaluate functional activities deferentially represented we mapped the DE genes to Gene Ontology (GO) terms (http://www.geneontology.org/). GO terms for the different ORFs were obtained from the PATRIC database (http://patricbrc.org/portal/portal/patric/Home). GO terms not present in the PATRIC database and whose annotation was obtained from the HOMD database or the J. Craig Venter Institute were acquired using the program blast2GO under the default settings ^6^. Enrichment analysis on these sets was performed using the R package ‘GOseq,’ which accounts for biases due to over-detection of long and highly expressed transcripts ^6^. Gene sets with ≤ ten genes were excluded from analysis. We used the REVIGO web page ^7^ to summarize and remove redundant GO terms. Only GO terms with FDR adjusted p-value < 0.05 in the ‘GOseq’ analysis were used.

### Phylogenetic assignment of transcripts (LEfSe)

Counts from the mRNA libraries were used to determine their phylogenetic composition for bacteria and archaea. Phylogenetic profiles of the metatranscriptomes were obtained using Kraken ^8^. We generated a custom Kraken library with the oral microbiome genomes indicated in the above section with a filtering threshold of 0.05. Phylogenetic profiles were used to identify significant differences between active communities under the different conditions studied by performing linear discriminant analysis (LDA) effect size (LEfSe) as proposed by Segata et al. ^9^ with an alpha value for the Wilcoxon test to 0.01. Significant *P*-values associated with microbial clades and functions identified by LEfSe were corrected for multiple hypothesis testing using the Benjamini and Hochberg false discovery rate correction ^10^ using the p.adjust function in R with a cutoff of FDR < 0.05.

